# A cell-topography based mechanism for ligand discrimination by the T-cell receptor

**DOI:** 10.1101/109785

**Authors:** Ricardo A. Fernandes, Kristina A. Ganzinger, Justin Tzou, Peter Jönsson, Steven F. Lee, Matthieu Palayret, Ana Mafalda Santos, Alexander R. Carr, Aleks Ponjavic, Veronica T. Chang, Charlotte Macleod, B. Christoffer Lagerholm, Alan E. Lindsay, Omer Dushek, Andreas Tilevik, Simon J. Davis, David Klenerman

## Abstract

The T-cell receptor (TCR) triggers the elimination of pathogens and tumors by T lymphocytes. In order for this to avoid damage to the host, the receptor has to discriminate between thousands of peptide ligands presented by each host cell. Exactly how the TCR does this is unknown. In resting T-cells, the TCR is largely unphosphorylated due to the dominance of phosphatases over kinases expressed at the cell surface. When agonist peptides are presented to the TCR by major histocompatibility complex (MHC) proteins expressed by antigen-presenting cells (APCs), very fast receptor triggering occurs, leading to TCR phosphorylation. Recent work suggests that this depends on the local exclusion of the phosphatases from regions of contact of the T cells with the APCs. Here, we develop and test a quantitative treatment of receptor triggering reliant only upon TCR dwell-time in phosphatase-depleted cell-cell contacts constrained in area by cell topography. Using the model and experimentally-derived parameters, we find that ligand discrimination is possible but that it depends crucially on individual contacts being 400 nm in diameter or smaller, *i.e.* the size generated by microvilli. The model not only correctly predicts the relative signaling potencies of known agonists and non-agonists, but achieves this in the absence of conventional, multi-step kinetic proof-reading. Our work provides a simple, quantitative and predictive molecular framework for understanding why TCR triggering is so selective and fast, and reveals that for some receptors, cell topography crucially influences signaling outcomes.

**Significance statement:** One approach to testing biological theories is to determine if they are predictive. A simple, theoretical treatment of TCR triggering suggests that ligand discrimination by the receptor relies on just two physical principles: (1) the time TCRs spend in cell-cell contacts depleted of large tyrosine phosphatases; and (2) constraints on contact size imposed by T cells using finger-like protrusions to interrogate their targets. The theory not only allows agonistic and non-agonistic TCR ligands to be distinguished but predicts the relative signalling potencies of agonists with remarkable accuracy. This suggests that the theory captures the essential features of receptor triggering.

## Introduction

T cells play a central role in immunity. Triggering of T-cell receptors (TCRs), expressed on the surfaces of all T cells, upon binding their ligands, peptide-MHC complexes (pMHC) present on antigen-presenting cells (APCs), sets T cells on course to respond to pathogens and tumors (1). The TCR’s capacity to distinguish between different MHC-bound peptide complexes is referred to as ligand discrimination, a process that crucially underpins immunological self/non-self recognition and T-cell development (2). Ineffective ligand discrimination often leads to immunodeficiency or autoimmunity (3). Despite its central role in immunity, the biophysical basis of ligand discrimination by the TCR is unclear, and understanding it is increasingly becoming a matter of considerable clinical significance. Engineered immune cells expressing re-purposed or artificial antigen receptors comprise a powerful new class of cancer therapeutics (4, 5). The severe off-target activity and extreme toxicity observed in some instances (6–8), however, is at least partly reflective of our poor grasp of the interplay between TCR binding kinetics, ligand density, and discriminatory signaling.

In addition to being highly selective, TCR signaling is extremely sensitive and fast: binding to a single agonist pMHC is sufficient to induce TCR signaling within seconds (9, 10). However, agonist peptides often comprise a very small fraction of all the peptides presented by APCs, raising the issue of how high sensitivity and discrimination are achieved simultaneously (11, 12). Several attempts have been made to explain ligand discrimination based on the TCR acting autonomously in ways analogous to G protein-coupled and growth factor receptors, with limited success. In general, TCR-induced signaling is assumed to rely exclusively on pMHC binding and thus, in modeling the process, little consideration is given to extrinsic factors that could influence signaling outcomes. Kinetic proofreading-based models of TCR signaling generally assume that a series of fast and slow downstream signaling steps allows ligand discrimination (13, 14), but the requirement for multiple steps comes at a cost: reduced sensitivity (15). Exactly how the compromise between ligand discrimination and sensitivity is reached remains poorly understood.

Triggering of the TCR results in the tyrosine phosphorylation of its intracellular signaling motifs by the kinase Lck, which initiates a cascade of chemical reactions in the T cell that leads to its activation. It has recently been shown that the large receptor-type tyrosine phosphatase CD45 is excluded from sites of close contact of T cells with model surfaces (16, 17), locally altering the kinase/phosphatase ratio in favor of ITAM phosphorylation (18). This raised the possibility that contact topography, *i.e.* cell shape, could have a significant bearing on signaling outcomes. When the phosphatase-excluding contacts formed with these surfaces are sufficiently large, signaling is spontaneously initiated even when TCR ligands are absent (16). But this in turn presents a paradox: whereas the TCR must discriminate between self and non-self ligands, *i.e.* between agonists and non-agonists, ligand binding does not appear even to be an essential feature of the triggering mechanism. These considerations suggest that the TCR cannot, and so does not, act autonomously to discriminate between ligands. Here, we develop and test a quantitative treatment of receptor signaling that relies only on TCR dwell-time at topographically constrained, phosphatase-depleted contacts, and explains ligand discrimination by the TCR, to the extent that the relative potencies of well-characterized pMHC ligands can be predicted with great accuracy.

## Results

### A signaling theory relying on TCR dwell-time at close contacts

The idea that TCR signalling might only depend on TCR ‘dwell-time’ at phosphatase-depleted regions of close contact between T cells and APCs is embodied in the kinetic-segregation (KS) model of TCR triggering (19). This model proposes that, at such contacts, the TCR remains accessible to kinases but is protected from phosphatases that would otherwise reverse its phosphorylation, resulting in the phosphorylated state being sufficiently long-lived for downstream signaling to be initiated. In this context, cognate pMHC ligands would promote signaling simply by increasing the dwell time of TCRs inside the close contacts, in part due to reduced diffusion as observed when TCRs interact with cognate pMHC presented on model surfaces (20, 21), but also because the TCRs become trapped in the contacts by ligand engagement.

In building a new quantitative treatment of signaling, we assumed only: *(i)* that when a close contact is formed, CD45 is excluded from the contact (**Fig. 1 *A***), *(ii)* TCRs can diffuse in and out of the contact, and *(iii)* that if the TCR is bound to a pMHC ligand there, it continues diffusing but is unable to leave the contact. Any TCR that remains in a close contact for longer than a minimal time *t_min_*, irrespective of ligand binding, is assumed to be triggered, *i.e.* a receptor ITAM is stably phosphorylated (**Fig. 1 *B***). *t_min_* is assumed to be 2 seconds, consistent with observation (10, 22–25). The value for effective Lck *k_cat_* that was used corresponded to that measured by Hui *et al.* (16) at the CD45/Lck ratio observed (18) in close contacts. *t_min_* thus creates an abrupt lower threshold for productive residence times. In addition to *t_min_*, the model incorporates the following parameters: *(i)* TCR density, *(ii)* TCR diffusion inside regions of close contact, which is greatly reduced by ligand binding, and *(iii)* the size and duration of the contacts, thereby explicitly including T-cell topography and dynamics. For freely diffusing TCRs in circular close contacts, the mean dwell-time (*τ_TCR_*) is dependent on contact radius *r* and inversely proportional to the diffusion coefficient *D* of the receptor:

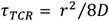

**Fig. 1.**
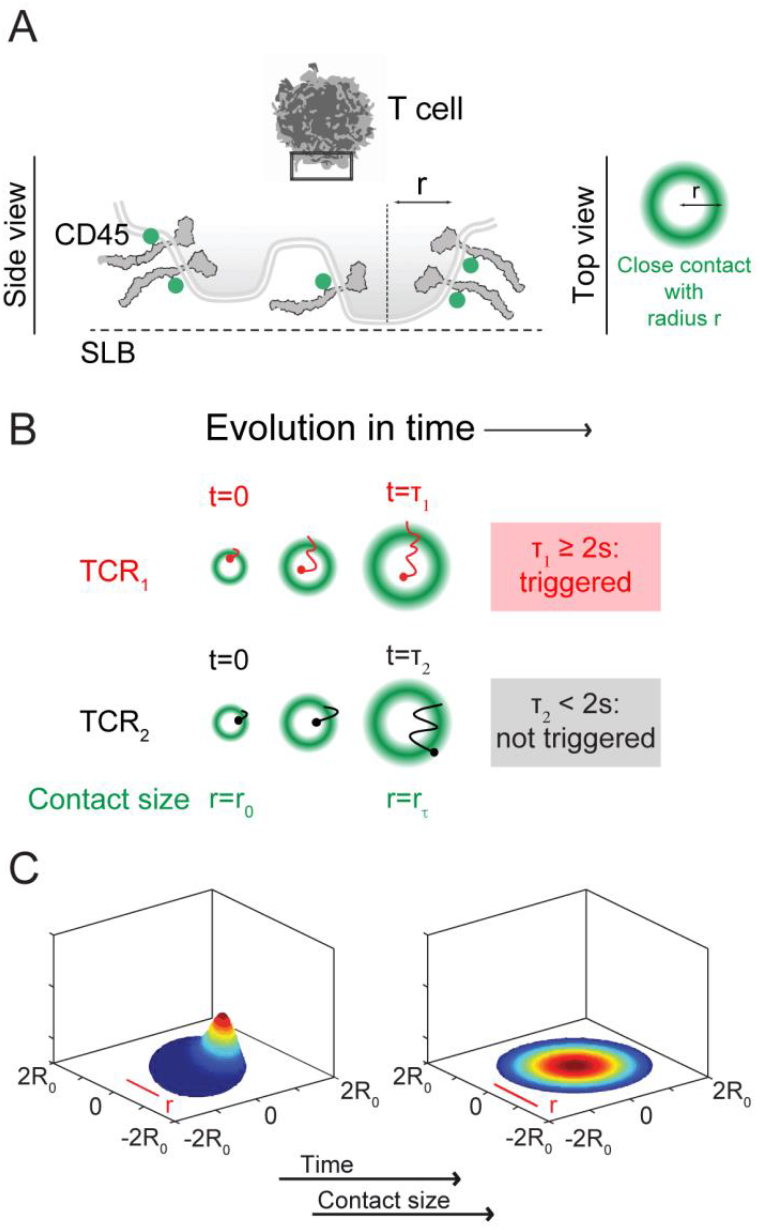
A quantitative treatment of TCR triggering reliant on receptor dwell-time inside close contacts. Top- and side-view depictions of main elements of the mathematical modelling: CD45 segregation and contact topography (area and duration) (*A*), and TCR mean residence time inside the close contact (*B*). Receptor triggering is assumed to occur for receptors that remain inside close contacts longer than 2 s. (*C*) shows snap shots from the simulation of the TCR density probability evolution in close-contacts as they grow over time.

However, because close contacts are not static and instead increase in area over time (16, 17), we calculated the probability that TCRs would dwell inside contacts for contacts that grow to radius *r*, assuming a radius-dependent rate of entrance of TCRs into the contacts (**Fig. 1 C, Movie S1**), by incorporating a moving-boundary analysis (for further details, see **Supplementary Information**: *Quantitative Treatment of Signaling*). Using this quantitative framework, we asked: what are the binding properties of ligands that would lead to TCR signaling? And what effect would contact size have? Most importantly, using the known binding and signaling properties of well-characterised ligands, we tested whether the model was predictive.

### Parameterization of the model

We parameterized the model by characterizing the features of T-cell interactions with supported lipid bilayers (SLBs), in the absence of TCR-ligands but with the defined membrane separation expected to be created *in vivo* by small adhesion receptors. For this, we used signaling-disabled forms of the rat adhesion proteins CD2/CD48, avoiding confounding effects introduced by adhesion receptor signaling *per se* (16, 26). Jurkat T-cells expressing signaling-disabled CD48-T92A (27) were allowed to settle onto SLBs presenting rCD2. We used two-colour total internal reflection fluorescence (TIRF) microscopy and single-molecule tracking to follow individual Lck, TCR or CD45 molecules (sub-stoichiometrically labelled) relative to the boundaries of close contacts identified by the presence of CD45 (labelled at densities above single-molecule resolution in a second color; **Fig. 2 *A***). CD45 exhibited the most exclusion from rCD2-mediated T-cell/SLB contacts. The density of CD45 molecules inside the close contacts was only 13±3% of that outside (**Fig. 2 *B*; Table S1**), versus 56±7% and 40±6% for Lck and the TCR, respectively (**Fig. 2 *C* and 2 *D*; Table S1**). The initial CD45/Lck ratio of 5 to 1 prior to contact was in this way reduced by ~50% (**Fig. S1**). The effective Lck activity at this CD45/Lck ratio has been shown to be ~half-maximal (close to 2.2 pTyr/s; (18)). Mean diffusion coefficients were similar for molecules inside and outside the contacts, and TCR diffusion was ~2-fold slower compared to CD45 and Lck (**Table S1** and **Fig. S2**).

**Fig. 2.**
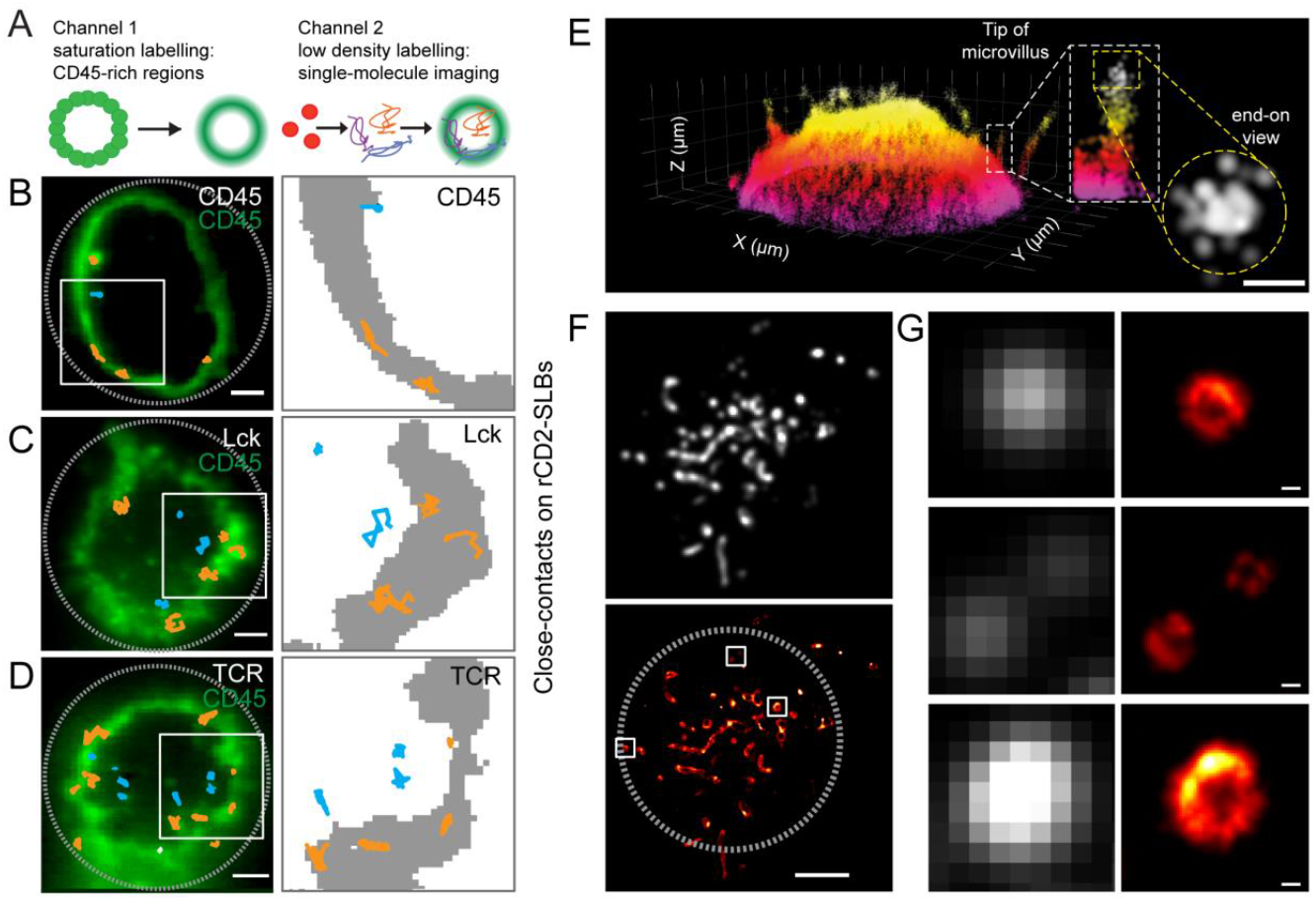
Single-molecule tracking of CD45, Lck and TCR distribution at close contacts formed with a model surface. (*A*) Experimental approach. High-density labeling of CD45 was used to indicate sites of close-contact formation between T cells and a rat CD2-presenting SLB *(left panel)*, and combined with simultaneous low-density labeling of CD45, Lck or TCR *(right panel)* to enable TIRF-based single-molecule tracking. *(B-D) Left panels:* TIRFM-based single-molecule tracking of CD45 (*B*), Lck (*C*) and TCR (*D*). Well-separated individual trajectories were recorded for >280 ms and coloured according to position in the contact (orange in CD45-rich region and blue in CD45-depleted region). *Right panels*: close-up views of trajectories in regions marked by white rectangles; CD45-rich regions are shown in grey. Scale bar, 2 μm. Data is representative of three independent experiments. (*E*) 3D super-resolution imaging (28) of fluorescently labelled CD45 in the Jurkat T-cell membrane *(left)* and side and tip of a microvillus (250 nm thick section, end-on view, *right);* localisation precision is ~15 nm laterally and ~30 nm axially. (*F*) 2D dSTORM image of CD45 in early T-cell contacts formed with IgG-coated glass; average *(top)* and reconstructed images (*bottom;* scale bar, 3 μm) are shown. *(G)* Close-up view of individual close-contacts (scale bar 100 nm). (*F*) and (*G*) are representative of three experiments.

Two assumptions of the model are *(i)* that CD45 is evenly distributed at the T-cell surface prior to contact formation, and *(ii)* that it is excluded as soon as close contacts are formed. 2D and 3D super-resolution imaging (28) confirmed that CD45 is indeed evenly distributed on the surface of T-cell microvilli (**Fig. 2 E**), consistent with previous findings (29), and that CD45 is spontaneously and passively excluded from contacts smaller in size than the diffraction limit within seconds of contact formation (~80 nm; **Fig. 2 *F, G;* Fig. S3 *A, B***; (17)). For Jurkat-CD48TM T-cells interacting with SLBs loaded with fluorescently-labeled CD45RABC and rCD2, the SLB-bound CD45 was spontaneously excluded from contacts that formed (**Fig. S3 *C*** and **Movie S2**), further confirming that CD45 segregation occurs passively. A summary of the key measurements made in this study or by others that were used for the modeling is given in **Table 1**; a more detailed list of parameters is presented in **Table S2**).

**Table 1.** Experimental parameters used in this study. References are given for measurements taken from the literature.

| Parameter |  |
| --- | --- |
| Total cell area | 415 [$\mu\text{m}^2$]<sup>1</sup> |
| Diffusion coefficient (TCR) | 0.05 [$\mu\text{m}^2\text{s}^{-1}$]<sup>2</sup> |
| Total # TCR per cell | 41,500<sup>3</sup> |
| Fraction of TCR segregation | 0.6<sup>2</sup> |
| CD45:Lck ratio | 2.5:1<sup>2</sup> |
| Close contact radius | variable |
| Mean cell contact diameter | 440 nm<sup>4</sup> |
| Half-life of Tcell-APC contacts | < 120 s<sup>4</sup> |
| Microvilli contact half times (no agonist) | variable |
| $t_{1/2}$ for weak "antagonist" | < 2 s<sup>4,5</sup> |

<sup>1</sup>Weaver, 1983
<sup>2</sup>Experimentally determined in this study for Jurkat T cells
<sup>3</sup>Experimentally determined in this study for CD4 T cells
<sup>4</sup>Cai et al, Science, 2017
<sup>5</sup>Hui and Vale, Nat Struct Mol Biol, 2014

### Validation of the model

The model calculates the probability that a TCR will remain in a close contact long enough for it to be triggered *(i.e.* phosphorylated), that is, longer than 2 seconds. It makes a number of testable predictions that we used for validation. Since the time, *τ_TCR_*, that the TCR must reside inside a close contact in order to be phosphorylated is dependent on the radius, which is modulated by the close-contact growth rate (if contacts grow on similar time scales to TCR diffusion), our model predicts that triggering times, that is, the time needed for at least one TCR to remain inside a close contact for longer than 2 seconds, will be shorter for cells with faster close contact growth-rates (*prediction 1;* **Fig. 3 A, *B***; see **Supplementary Information** for further details). In addition, because the phosphorylation rate, *i.e.* the effective *k_cat_* of Lck, changes according to the CD45/Lck ratio, an increase in this ratio should lead to an increased t_min_, requiring longer TCR dwell-times for a TCR to be triggered (*i.e.* longer triggering times would be observed, *prediction 2;* **Fig. 3 *B***; for quantification of Lck *k*_cat_ at different CD45/Lck ratios see ref 18). Finally, receptor triggering is predicted to occur sooner for single large contacts compared to two separate contacts of the same combined size (*prediction 3*). For example, the model predicts that the triggering probability increases by more than 7-fold when two single contacts coalesce into a larger one (**Fig. 3 *C*; see also Supplementary Information**).

**Fig. 3.**
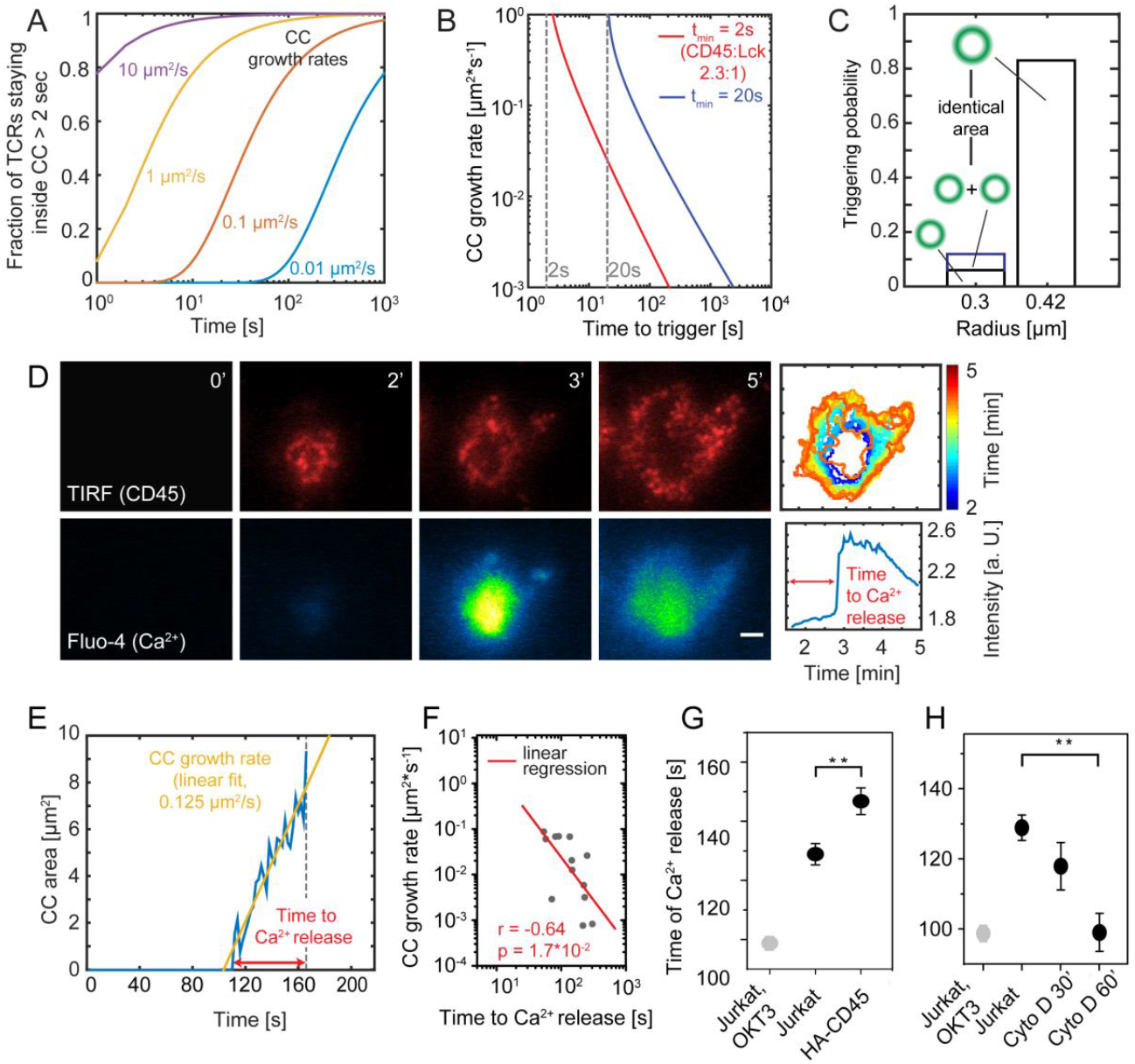
Tests of the model indicate that TCR signalling can be explained by TCR dwell-time in close contacts. (*A*) Fraction of triggered TCRs as a function of time and contact growth-rate (t_min_ = 2 s, D = 0.05 μm^2^/s, g = 0.01 – 10 μm^2^/s). (*B*) Time taken to TCR triggering as a function of close contact growth-rate. (*C*) Comparison of triggering probability for one versus two contacts or a single contact of double the contact area. (*D*) Dynamics of close-contact formation (CD45 fluorescence, TIRFM; *top panels*) and Ca^2+^ release (detected as Fluo-4 fluorescence, *bottom panels*) for cells contacting rCD2-presenting SLBs. Scale bar, 2 μm. *Top right panel:* color-coded representation of the temporal evolution of contact area over time. *Bottom right panel:* temporal evolution of Fluo-4 intensity averaged over entire contact. (*E*) Trace of a representative contact over time for growth-rate determination. (*F*) Relationship between close contact growth-rate and the time taken to triggering. *(G)* Time delay between initial contact of cells with rCD2-presenting SLBs and Ca^2+^ release for Jurkat T-cells and cells expressing HA-CD45. *(H)* Time delay between initial contact of cells with IgG-coated glass and Ca^2+^ release in the presence of the actin depolymerising drug cytochalasin D (**p=0.01 and <0.001, two-tailed t-test, unequal variance assumed; errors are s.e.m.).

To validate the model, we set out to test these predictions quantitatively, using calcium release as a proxy for ITAM phosphorylation. To test *prediction 1*, we simultaneously measured close contact growth-rates and signaling times by coupling TIRF-based identification of close contacts formed by T cells interacting with rCD2-presenting SLBs, with signaling detected as calcium release (**Fig. 3 *D, E*** and **Movie S3**). Contact formation was detected *via* the exclusion of CD45 at sites of interaction with the surface (16). In agreement with the model’s prediction, receptor triggering occurred faster in cells with larger close contact growth-rates (**Fig. 3 F**). To test *prediction 2*, we compared the triggering times for wild-type Jurkat T-cells with those for cells expressing a form of CD45 (HA-CD45) lacking its extracellular domain, which is less efficiently excluded from contacts, thereby reducing Lck k_cat_ by increasing the CD45/Lck ratio in the close contacts (**Fig. S1**; *(10)).* As predicted once again by the model, expression of HA-CD45 at ~10,000 copies/cell *(i.e.* at 5% of total CD45 expression; **Fig. S5**) delayed triggering by almost 20 s (~15%, p < 0.05, two-tailed t test, unequal variance assumed; **Fig. 4 *G***). Finally, treatment of Jurkat T-cells with cytochalasin D, an inhibitor of actin polymerization and microvillus formation (30) that produced larger and more stable contacts, led to faster triggering times by up to 30 s (~23%, p < 0.05) in a drug exposure-dependent manner, consistent with the third prediction of the model (**Fig. 4 *H***).

**Fig. 4.**
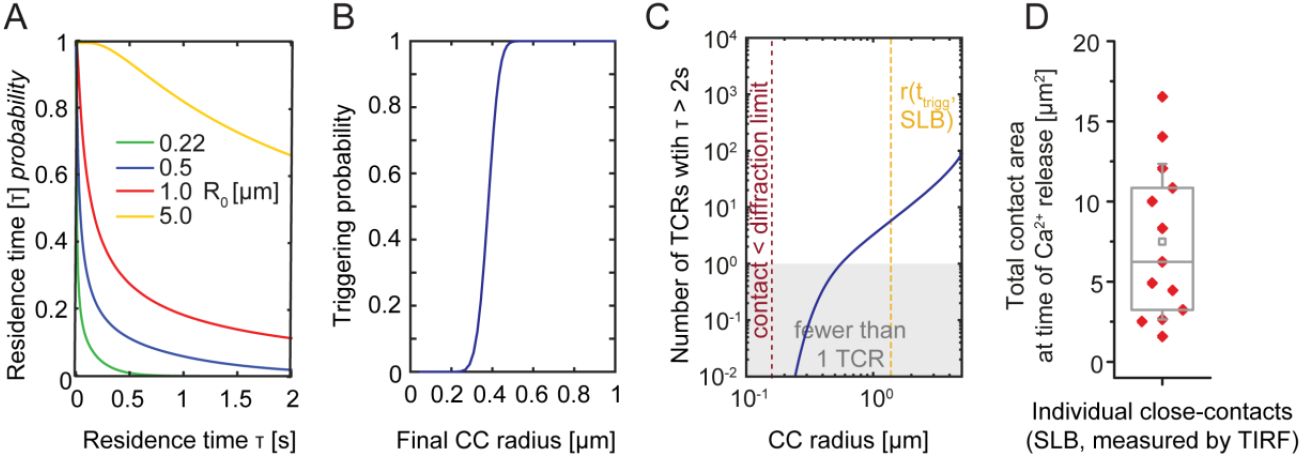
Calculations of TCR triggering probability. *(A)* Probability that a TCR remains inside a close contact for time τ, for close contacts of varying fixed radius, R_0_. *(B)* Probability that a single TCR stays inside a close contact > 2 s as a function of final close-contact radius for growing contacts. *(C)* Total number of TCRs that remain inside the close contact for > 2 seconds, incorporating the estimates shown in (*A*), the density of TCRs in Jurkat T cells, and the degree of exclusion of the TCR from close contacts for cells interacting with rCD2-presenting SLBs. *(D)* Total contact area (area of CD45 exclusion) at the time of calcium release for T cells interacting with rCD2-presenting SLBs (13 cells). Central lines indicate the median; small squares indicate the mean; boxes show interquartile range; whiskers, s.d.

### Signaling in the absence of ligands

Having validated the model, we explored its implications for signaling. We calculated the dwell-times of individual TCRs in close contacts and determined the probability *p* that any given receptor resides there for *t >t_min_ = 2 s*, which we refer to as “triggering probability”. We found that *p* increases sharply with increasing close contact radius (**Fig. 4 *A, B***). In the absence of ligands, *p* is ~0 for contacts of the size observed during T-cell interrogation of APCs, ~0.22 μm (**Fig. 4 A**; (31)), implying that no TCR is likely to be triggered in such contacts. On SLBs, T cells form contacts (**Fig. 2 *B-D***) much larger than those observed during cell-cell interactions, and for these types of contacts we estimate that ~16 TCRs will be triggered per contact in the absence of ligands (**Fig. 4 C**; the grey area corresponds to fewer than one TCR staying in the contact for *t_min_* of at least 2 s). We determined this value using *(i)* triggering probability *p, (ii)* the total contact size observed at the time of calcium signaling (median contact area of 6 μm^2^; **Fig. 4 *D***), *(iii)* the measured overall TCR density (**Fig. S4**), and *(iv)* the fraction of TCRs inside the contacts (40%; **Fig. 2 D**). Previously we have shown that TCR triggering in the absence of ligands is readily observable under these conditions (16), in good agreement with the model’s predictions.

For close contacts of the size observed when T cells encounter antigen-presenting cells *(i.e.* r= 0. 22 μm; **Table 1**; (31, 32)), it takes ~18 hours to reach a 50% probability that a given TCR is triggered without ligands (**Fig. 5 *A*** and **Supplementary Information**). Strikingly, however, only a 2-fold increase in close contact radius yielded a ~1,000-fold increase in the probability of TCR triggering in the absence of ligands (50% signaling within 70 s; **Fig. 5 *A***). This suggests that the size of the contacts used by T-cells to interrogate APCs is close to, but just below the threshold size that would produce non-specific, ligand-independent TCR triggering. For microvillar contacts with half-lives similar to those observed in the absence of ligands (31), TCR dwell times of even 1 s, *i.e.* half of t_min_, are highly unlikely (**Fig. 5 *B***). These observations suggest that ligand dependent T-cell responses are highly reliant on contact size being constrained, and that signaling without ligands is further avoided by microvillar contacts being short-lived.

**Fig. 5.**
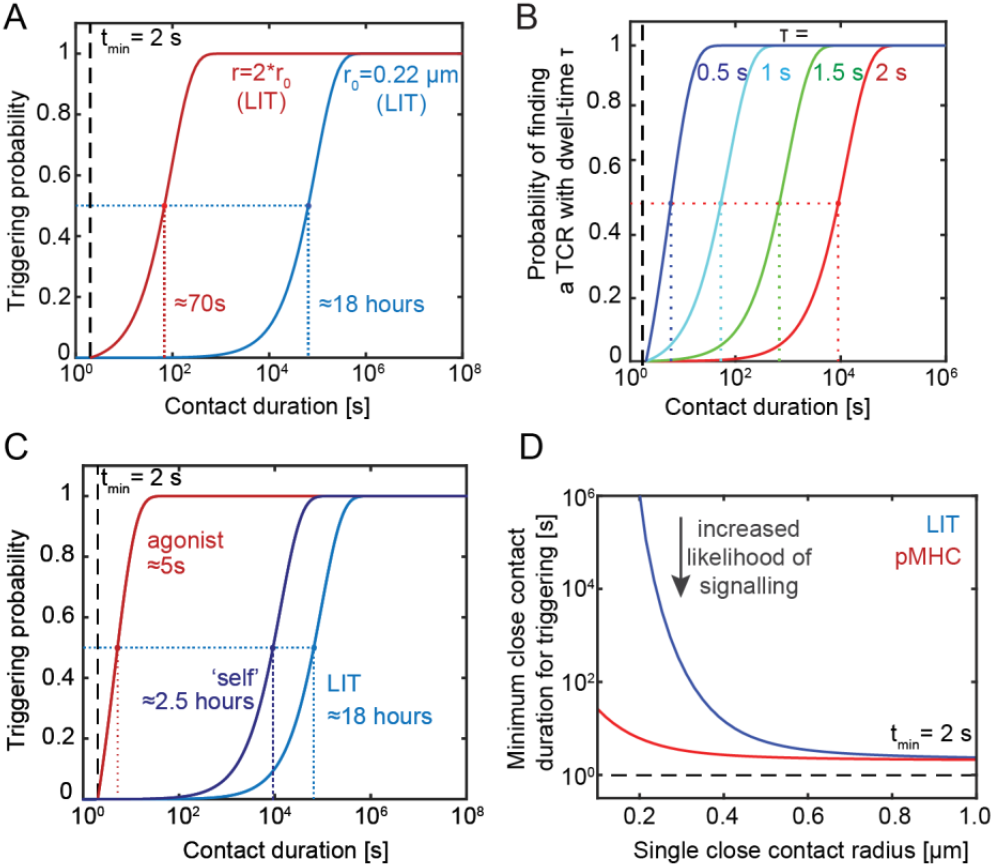
Dependence of TCR triggering on ligand-binding and close contact size. (*A*) Probability that at least one TCR will be triggered (*i.e.* stay in the contact > 2s), as a function of contact duration in the absence of ligand. Triggering probability is calculated for a single close-contact of 0.22 μm radius and for a larger close-contact of 0.44 μm (LIT, ligand-independent TCR triggering). (*B*) Probabilities of finding TCRs with dwell-times of 0.5, 1, 1.5 and 2 s in the absence of ligand. (*C*) Probability that at least one TCR will trigger as a function of contact duration in the presence and absence of agonist pMHC with a low *k*_off_ (*k*_off_ = 1, 30 pMHC molecules/μm^2^), and of a pMHC with a larger *k*_off_ at higher pMHC densities (*k*_off_ = 50, 300 pMHC molecules/μm^2^), for a close contact of 0.22 μm radius. (D) Effect of close contact radius on the contact duration required for TCR triggering in the presence or absence of agonist pMHC (*k*_off_ = 1; 30 pMHC molecules/μm^2^).

### Self/non-self discrimination

We next addressed the problem of discrimination (see **Supplementary Information** and **Table S2** for further details). Agonist pMHC, encountered at 0.22 μm contacts even at low density (30 pMHC/μm^2^2D K_d_ given by *k*_off_ = 1 ^−1^ and *k*_on_ = 0.1 μm^2^s^−1^), increased the triggering probability by ~12,000-fold, *i.e.* from 18 hours to 5 seconds (**Fig. 5 *C***). For pMHC/TCR interactions with *k*_off_ = 50 s^−1^ and for pMHC at 300 molecules/μm^2^, *i.e.* at the observed low-affinity threshold for non-agonistic TCR/pMHC interactions at high ligand-density (2, 11, 33, 34), 50% TCR triggering probability still required stable contacts lasting 2.5 hours (**Fig. 5 *C***). In other words, a 50-fold increase in *k*_off_, reflecting a very conservative estimate of the lower limit of the *k*_off_ for self pMHC, led to a 1,800-fold reduction in the likelihood of TCR triggering occurring, despite there being 10-fold more ‘self’ binding molecules than agonist pMHCs. As expected, changes to either *k*_on_, *k*_off_ or pMHC density altered the triggering probability profoundly (**Fig. S6 *A***). Whereas ligand-independent TCR signaling was markedly dependent on close-contact radius, agonist signaling was not, suggesting that contact topology has the capacity to skew signaling in favor of agonists (**Fig. 5 *D***).

Kinetic proof-reading (KP) schemes are typically used to explain discriminatory signaling by the TCR. Classically, KP requires multiple intermediate steps to produce distinct signaling outcomes for ligands of different affinity and potency (13). In these types of calculations, for example, six intermediate steps allows a 10-fold difference in pMHC affinity to generate a >7,500-fold difference in TCR triggering. Such large amplification mechanisms are only possible, however, at the expense of sensitivity (13, 15). In the new model, the same 10-fold affinity difference results in a >1,800-fold difference in TCR signaling for pMHC densities of 10-30 molecules/μm^2^ (**Fig. 6 *A***). Even at very low pMHC densities (1 molecule/μm^2^), there was a >300-fold difference in TCR signaling probability. This high level of discrimination also occurs very rapidly, since the TCR becomes phosphorylated in a single step taking 2 seconds. In contrast to classic KP schemes, therefore, the new approach allows for rapid, high-level discrimination even at low pMHC densities.

**Fig. 6.**
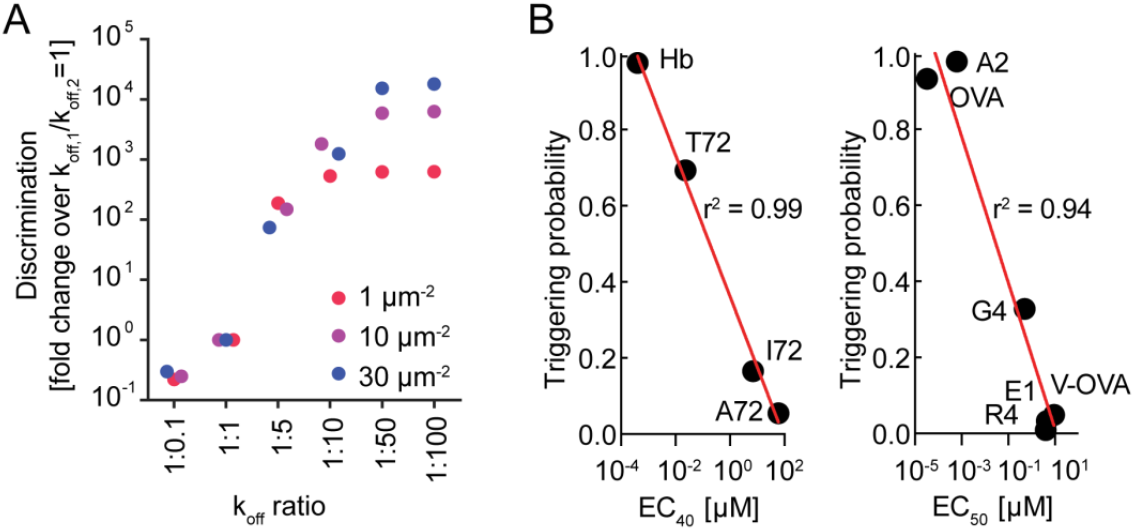
Predictions of T-cell signalling probability based on known 2D TCR/pMHC binding properties alone. (*A*) Analysis of discrimination (relative triggering probabilities) for pMHC ligands forming TCR complexes with different off-rates, and present at different densities. (*B*) Peptide stimulation potencies (EC_40_ and EC_50_ values for IL-2 secretion) for CD4^+^ *(left panel)* and CD8^+^ T-cells *(right panel;* determined elsewhere (11, 36)), plotted against the probability that at least one TCR triggering event (τ > 2 s) occurs at a single contact of 0.22 μm radius that persists for 120s (see **Table S3** for further details).

Finally, we tested whether we could predict the potency of TCR ligands at cell-cell contacts, relying only on experimentally-determined 2D *k*on and *k*off values (the parameters used are given in **Table 1 and Table S2**). For ten different pMHC complexes, the calculated TCR triggering probability was found to correlate remarkably well with signaling potency, measured as IL-2 production in cocultures of peptide-pulsed APCs and T cells (**Fig. 6 B** and **Table S3**) (11, 35, 36).

## Discussion

Prompted by the finding that phosphatase exclusion at close contacts formed by T cells interacting with model surfaces suffices to induce TCR triggering (16), we have explored the implications of TCR signaling depending only on receptor dwell-time in phosphatase-depleted contacts. Using a new quantitative treatment of TCR signaling, we find that we can explain both ligand discrimination and sensitivity.

The model used measurements of *(i)* CD45/Lck ratio at close contacts; *(ii)* TCR density and diffusion; *(iii)* the size and duration of close contacts; and *(iv)* estimates of Lck activity expected at the levels of CD45/Lck segregation observed at the contacts. Validating the model, we showed that close-contact growth-rate and triggering time were inversely correlated, that signaling was delayed by lower CD45 segregation, and faster when contact area was increased. The probability of ligand-independent TCR triggering was found to increase sharply for close contacts of radius larger than ~0.2 μm, close to the dimensions of contacts observed *in vivo* (29, 31, 32, 37). For the much larger close contacts formed on SLBs in the absence of ligands (r~0.65 μm), we calculated that ~80 TCRs would be present and that ~20% of these would be triggered. Similar numbers of TCRs have to be engaged by conventional ligands to observe signaling when CD4^+^ T-cells interact with APCs in the absence of co-receptors (~30 TCRs (10)). Overall, our observations suggest that TCR triggering might be so simple as to depend only on receptor occupancy of close contacts, with mean residence time having only to exceed a threshold for receptor triggering to occur. We propose that pMHC specific responses are governed by the kinetics of the TCR/pMHC interaction along with TCR diffusion, CD45 segregation and T-cell topography, since each of these will influence the mean residence time.

We asked how, in this context, the TCR discriminates between different-quality ligands. Without ligands and for close contacts of the size observed *in vivo, i.e. r=0.22* μm, a T-cell contact would need to persist for almost a day to trigger a single TCR. This implies that ligand-independent signaling, although easily demonstrated *in vitro*, is unlikely ever to occur *in vivo.* A surprising finding, however, was the remarkable sensitivity of ligand-independent signaling to close contact area. A 2-fold increase in contact radius enhanced signaling probability by three orders-of-magnitude, making it likely to occur within 70 s. The size of close contacts observed to form *in vivo* seems therefore to be close to the threshold needed to ensure that signaling is pMHC-dependent.

In contrast to most other receptors, such as GPCRs that are triggered in a largely binary fashion by single ligands, the TCR can react to multiple ligands varying up to 10^6^-fold in affinity (38). This level of discrimination is proposed to derive from kinetic proofreading (13). In a previous simulation of the kinetic-segregation model, multiple steps and long delays were required for effective proof-reading because kinase activity was assumed to increase 200-fold inside versus outside close contacts, resulting in even short-lived complexes being phosphorylated (39). Our calculations suggest, however, that discrimination is achievable in a single step: *i.e.* receptor phosphorylation. This is perhaps because the modest increases in net kinase activity inside close contacts greatly reduces the likelihood that weakly-bound receptors will be phosphorylated. An additional, theoretical possibility worth exploring is that limited sampling of self pMHCs at small contacts reduces the stochastic background.

Utilizing single step proof-reading, we could predict the relative potency of pMHC ligands with remarkable accuracy (r^2^=0.94-0.99). The previous best predictions were obtained by Aleksic *et al.* (r^2^=0.83), using the concept of ‘confinement time’ (the total time a TCR is occupied by pMHC before complete dissociation (40)). The improved predictive ability of the new model likely arises partly due to our use of 2D rather than 3D binding parameters, but mostly because of the spatial constraints imposed by limiting contact to a well-defined, small area. Our analysis also showed that the level of very early signaling (*i.e.* ITAM phosphorylation) predicts the scale of a late signaling outcome (IL-2 release). The striking relationship between agonist pMHC potency, TCR triggering and cytokine release has been noted before (9) and suggests that, at a single-cell level, T-cell activation is digital and perhaps wholly dependent on the probability of TCR triggering events.

Our findings have several important implications. First, the size of close contacts committing T cells to synapse formation may have to be tightly controlled to avoid non-specific activation. Defects in processes that constrain close contact size could predispose to autoimmunity by increasing ligand-independent receptor signaling. Second, we can estimate the extent to which the probability of TCR signaling is enhanced by pMHC binding, without the triggering mechanism having to be strictly ligand dependent. For TCRs interacting with typical ligands at small contacts, we calculated that agonist-dependent signaling is favored as much as 12,000-fold over ligand-independent signaling. Third, some degree of signaling in the absence of ligand might nevertheless explain both TCR polarization and partial TCR phosphorylation (41, 42). We estimate that ~50% of TCRs remain >0.5 s inside close contacts of 0.22 μm radius, yielding >1 pTyr/contact. This might not initiate downstream signaling but could generate the pMHC-independent, low-level “tonic” TCR signaling observed *in vivo* (41). Fourth, for close contacts increasing in radius beyond 0.22 μm, perhaps following an initial round of ligand-dependent signaling, ligand-independent receptor triggering might reinforce or amplify the initial response, enhancing sensitivity. Lastly, the principles established here could be extended to other ITAM-based receptors that are also sensitive to size-based changes in the kinase/CD45 ratio, such as Fc receptors (43), or used to calculate the binding ‘sweet-spot’ for engineered TCRs (4) or receptor mimics (44).

In conclusion, our work suggests that topographically-constrained T-cell contact formation is compatible with, and may even be essential for, ligand discrimination by T cells. The new model’s ability to predict the relative signaling potencies of known agonists and non-agonists suggests that it captures the essential features of the TCR triggering mechanism. But how might contacts be constrained? T-cell microvilli are the obvious candidates for achieving this. Microvillus-based contacts have radii of 0.22 ± 0.02 μm (32), and persist for 1-5 minutes (37, 45, 46). Individual microvillar contacts last >6 s in the absence of cognate antigen, enough time for efficient discriminatory signaling according to our calculations. Our new treatment of TCR signaling therefore provides a new, predictive framework for understanding why TCR triggering is selective, fast and sensitive, and explains why T cells interrogate their targets using microvilli.

## Acknowledgements

This work was funded by The Wellcome Trust, the UK Medical Research Council, the UK Biotechnology and Biological Sciences Research Council and Cancer Research UK. We thank the Wolfson Imaging Centre, University of Oxford, for access to the microscope facility. We would like to thank the Wellcome Trust for the Sir Henry Dale Fellowship of RAF (WT101609MA), the Royal Society for the University Research Fellowship of SFL (UF120277) and acknowledge a GSK Professorship (DK). We are also grateful to Doug Tischer (UCSF, US) and Muaz Rushdi (Georgia Tech, US) for their critical comments on the manuscript.

## Author contributions

R.A.F. and K.A.G. designed and performed experiments, analyzed data and R.A.F., K.A.G., S.J.D. and D.K. wrote the manuscript with input from all authors. J.T., A.E.L, O.D. and A.T., designed all mathematical modelling. P.J. co-performed some of the TIRF experiments and S.F.L., A.R.C., A.P. and M.P. super-resolution microscopy. A.M.S. performed calcium signaling experiments. V.T.C expressed and purified soluble CD45. B.C.M. designed various imaging setups. C.M. performed single molecule tracking experiments under supervision of K.A.G. S.J.D., A.T., and D.K. conceived and supervised the project.

## Supplementary Information

### Contents

**I. Online Materials and Methods**

**II. Supplementary Tables**

**Supplementary Table 1** | Quantitative characterization of close contacts between T cells and rCD2-SLB and the organization of key signaling proteins within these areas as determined by single-molecule fluorescence microscopy.

**Supplementary Table 2** | Parameters used for the mathematical modelling to derive the mathematical analysis represented throughout Fig. 3, 4, 5 and 6.

**Supplementary Table 3** | Detailed pMHC/TCR density and kinetic parameters used for the calculations shown in Fig. 6 B.

**III. Supplementary Figures**

**Fig. S1** | Quantitative analysis of CD45 and Lck density in close-contacts.

**Fig. S2** | TCR diffusion at close-contacts between T cells and supported lipid bilayers by single-particle tracking

**Fig. S3** | CD45 is equally distributed on resting T cell surfaces and segregates from close-contacts between T cells and glass surfaces at sub-μm length scales.

**Fig. S4** | Measurement of total TCR numbers in the cell line used for experiments.

**Fig. S5** | Measurement of HA-CD45 expression levels.

**Fig. S6** | The effect of TCR/pMHC complex life time on triggering probabilities and the contribution of LIT to the overall signalling probability at varying close-contact radii and for ligands with different k_off_ rates.

**IV. Supplementary Movies**

**Movie S1** | Animation of the changes in the TCR probability density across a growing close contact over time corresponding to the modelling shown in Fig. 1g.

**Movie S2** | CD45-RABC spontaneously segregates from CD2-mediated close-contacts formed by Jurkat T-cells (CD48+) with CD2- and CD45RABC-containing SLBs

**Movie S3** | Ca^2+^ release as measured by change in Fluo-4 fluorescence in Jurkat T-cells forming contacts depleted of CD45 (labeled) with rCD2-containing SLBs.

### Materials and Methods

#### Cell lines

Human Jurkat T cells (clone E6-1) and J.RT3-T3.5 were obtained from ATCC (ATCC^®^TIB-152™). All other cell lines were transduced with lentivirus to express gene of choice. Human HEK293T were obtained from ATCC (ATCC^®^ CRL-3216™). T Cells were cultured in sterile RPMI (Sigma Aldrich) supplemented with 10% FCS (PAA), 2 mM L-Glutamine (Sigma Aldrich), 1 mM Sodium Pyruvate (Sigma Aldrich), 10mM HEPES (Sigma Aldrich), and 1% Penicillin-Streptomycin-Neomycin solution (Sigma Aldrich). HEK293T cells were cultured in sterile DMEM (Sigma Aldrich) supplemented with 10% FCS (PAA), 2mM L-Glutamine (Sigma Aldrich) and 1% Penicillin-Streptomycin (Sigma Aldrich) at 37 °C and 5% CO_2_ and were maintained between 10% to 90% confluency. Cells were maintained at 37 °C and 5% CO_2_ during culturing, and handling was performed in HEPA-filtered microbiological safety cabinets. Typically, cells were kept at a density between 5-9 x 10^5^ cells/ml. The UCHT1 hybridoma was a generous gift from Dr Neil Barclay, Sir William Dunn School of Pathology, University of Oxford and the GAP8.3 hybridoma was obtained from ATCC (HB-12™). Hybridoma cells were cultured in sterile DMEM (Sigma Aldrich) supplemented with 10% FCS (PAA), 2mM L-Glutamine (Sigma Aldrich) and 1 mM Sodium Pyruvate (Sigma Aldrich) at 37 °C and 5% CO2 and were maintained between 10% to 90% confluency. All cells used throughout this study were regularly tested for mycoplasma.

#### Plasmids

For HA-CD45-Halo, LCK-Halo, and TCRβ-Halo (New England Biolabs, UK) the genes were amplified by PCR to produce dsDNA fragments encoding proteins of interest flanked at the 3’ end by a sequence coding for a Gly-Ser linker which was followed by Halo-tag. Following confirmation of sequence and reading frame integrity the Lck-Halo and TCRβ-Halo were sub-cloned into the lentiviral pHR-SIN plasmid. To generate mm-Lck the appropriate residues were mutated by a PCR amplification reaction using forward and reverse oligos encoding the desired mutation. Sequence integrity was confirmed by reversible terminator base sequencing.

#### Generation of stable transduced cell lines

Jurkat derived T cell lines stably expressing either mCitrine-actin, LCK-Halo, and TCRβ-Halo were generated using a lentiviral transduction strategy. HEK293T cells were plated in 6-well plates at 6 x 10^5^ cells per well in DMEM (Sigma Aldrich), 10% FCS (PAA) and antibiotics. Cells were incubated at 37 °C and 5% CO_2_ for 24h before transfection with 0.5 μg/well/plasmid of the lentiviral packaging vectors p8.91 and pMD.G (2^nd^ generation) and the relevant pHR-SIN lentiviral expression vector using GeneJuice^®^ (Merck Millipore) as per the manufacturer’s instructions. 48 h post transfection, the supernatant was harvested and filtered using a 0.45 μm Millex^®^-GP syringe filter unit to remove detached HEK293T cells. 3ml of the lentiviral-conditioned medium was added to 1.5 x 10^6^ Jurkat T cells.

#### Fab preparation and labeling

The TCR on Jurkat T cells was labeled with Alexa Fluor 647 Fab (UCHT1, anti-CD3ε; purified from hybridoma supernatant). The CD45 was labeled with Alexa Fluor 488 Fab (Gap8.3, anti-CD45; purified from hybridoma supernatant). Both Fabs were prepared from purified antibody using immobilized papain (agarose resin, ThermoFisher and per manufacturer protocol). Fab digestion and purity was confirmed by size exclusion chromatography. For Fab labeling, Alexa Fluor 488 and Alexa Fluor 647 antibody labeling kit (ThermoFisher) was used as per manufacturer protocol. For cell labeling, 1 ml of 5 x 10^5^ cells/ml were incubated with Fab (1-10 nM) on ice for 25 minutes. Cells were washed three times in 20 nm filtered PBS.

#### Fluorescence-activated cell sorting and quantification of protein expression

Wild type or transduced Jurkat and HEK293T cells were washed once in ice-cold PBS, and 1 million cells were incubated with appropriate antibodies (isotype control, eBioscience, UK; Gap 8.3, purified from hybridoma cells; anti-HA, clone HA-7, Sigma Aldrich; UCHT1, purified from hybridoma) at 10 ug/ml for 30 min on ice in PBS/0.05% Azide, washed once in PBS and incubated with a fluorescently labeled secondary anti-mouse antibody as appropriate (alexa 647 or alexa 488, Molecular Probes, Invitrogen or PE-conjugated, Sigma Aldrich) for a further 30 min on ice. Cells were washed in ice-cold PBS and analysed on a Beckman Coulter CyAn Analysers. For quantification of cell surface protein expression QuantiBrite-PE beads (BD Biosciences) were used as per manufacturer instructions. Mean fluoresce intensity analysis and further data processing was performed with FlowJo software using a standard gate on live cells based on the forward- and side-scatter profile.

#### HaloTag^®^ labelling

Cells expressing HaloTag^®^ (Promega, UK) fusion protein were labeled with TMR Cell * following manufacturer’s preparation protocol (www.promega.co.uk/products/imaging-and-immunological-detection/cellular-imagingwithhalotag/proteintrafficking). First, the cell medium was replaced with 200 μl RPMI without supplements to which 1-5 μM of Halo-Tag TMR dye was added and cells were incubated at 37°C for 45 minutes. To ensure that free dye would not remain in the cytoplasm, cells were washed three times in HBS and then further incubated 37°C for 30 minutes followed by another three washes with PBS.

#### Sample preparation for Calcium response measurements (Fig. 4 *D-H*)

Jurkat T cells were labeled with 4 μM Fluo-4 AM (F-14201; Invitrogen, Paisley UK) for 30 min at room temperature with 2.5 mM probenecid (P-36400; Invitrogen, Paisley UK) in RPMI (Sigma-Aldrich, UK) without supplements. Cells were then washed in HBS (51558; Sigma, UK) and the medium changed to HBS containing 2.5 mM probenecid before their addition to the microscope sample container with the prepared microscope coverslip.

#### Total internal reflection microscopy (TIRFM)

*For all Figures containing TIRF data (except Fig. 2 E-G):* TIRF imaging was performed using total internal reflection fluorescence microscopy (TIRFM). A diode laser operating at 488 nm (20mW, Spectra Physics, Newport, US) and either a diode laser operating at 561 nm (Excelsior, 20mW, Spectra Physics, Newport, US) or a HeNe laser operating at 633 nm (25LHP991230, Melles Griot) were directed into a TIRF objective (60x Plan Apo TIRF, NA 1.45, Nikon Corporation, Tokyo, Japan) mounted on an Eclipse TE2000-U microscope (Nikon Corporation, Tokyo, Japan) parallel to the optical axis and offset in order to achieve total internal reflection of the beam. The emitted fluorescence was collected by the same objective and separated from the returning TIR beam by a dichroic mirror (FF500/646-Di1, 488/633 emission, Semrock, US) or (XF2044-490-575DBDR, 488/561 emission, Omega Optics). The green fluorescence emission was subsequently separated from the red fluorescence emission by a second dichroic mirror and filter sets; for excitation with 488 and 633: FF605-Di02 (Dual-View mounted, Photometrics, Roper Scientifics, US), FF03-525/50-25 (488 emission), BLP01-635R-25 (633 emission), all Semrock, US; for excitation with 488 and 561: FF562-Di03 (Dual-View mounted, Photometrics, Roper Scientifics), FF02-525/40-25 (488 emission), LP02-568RS-25 (561 emission), all Semrock, US. The fluorescence signals from both channels were simultaneously recorded using an EM-CCD camera (Cascade II: 512, Photometrics, Roper Scientifics, US) operating at −70 °C, whereby each color was recorded on one half of the EMCCD chip. A grid consisting of regularly spaced ion-beam etched holes in gold-on-glass was used to achieve image registration across both emission channels. The Dual-View optics were adjusted to maximize the overlap of the grid images in the two channels under bright field illumination, resulting in a mean alignment precision of approximately 120 nm. Data were acquired using either single snap shots or time-lapse acquisition using Micromanager (Edelstein et al., 2010).

*For Fig. 2 F,G:* A collimated 640 nm (LaserBoxx 641, Oxxius, Lannion, France) laser beam was focused at the back aperture of a 60x oil TIRF objective (Olympus, NA 1.49, UIS2 series APON 60XOTIRF) mounted on an IX71 Olympus inverted microscope frame. The power of the collimated beams at the back aperture of the microscope was 20 mW. Emitted fluorescence was collected by the same objective, separated from the excitation by a dichroic (Di01-R4-5/488/561/635, Semrock), expanded through a 2.5x achromatic beam expander (Olympus, PE 2.5x 125) and sent through an emission filters 635 long-pass filter (Semrock, BLP01-635R). The images were recorded on an EM-CCD camera (Evolve 512, Photometrics) operating in frame transfer mode at −80°C. Videos of 1000 frames were acquired at exposure times of 30 ms using Micromanager.

#### Double-helix Point-spread function (DHPSF) microscopy (Fig. 2 E)

As previously described in (Carr et al., 2017), DHPSF imaging was carried out on a bespoke microscope incorporating a 1.27 NA water immersion objective lens (CFI Plan Apo IR SR 60XWI, Nikon) mounted on a scanning piezo stage (P-726 PIFOC, PI) onto a conventional fluorescence microscope body (Eclipse Ti-U, Nikon). A 4f system of lenses with a 650 nm optimised double-helix phase mask (DoubleHelix, Boulder USA) placed in the Fourier-plane performed the DHPSF transformation and relayed the image onto an EMCCD detector (Evolve Delta 512, Photometrics). Collimated 640 nm (200 mW, iBeam smart-640-s, Toptica) and 405 nm (120 mW, iBeam smart-405-s, Toptica) lasers were directed into the objective lens, resulting in a power density at the sample of ~1 kW/cm^2^ and 100 W/cm^2^ respectively. The fluorescence signal was separated from the laser excitation by a quad-band dichroic filter (Di01-R405/488/561/635-25x36, Semrock). Additional isolation of the fluorescence signal was achieved by long-pass and band-pass filterers placed directly before the detector.

#### Supported Lipid Bilayers (SLB) preparation

Prior to SLB formation on glass cover slips, the cover slips (size no. 1: 0.13 mm in thickness, VWR International, UK) were cleaned by incubation in Piranha solution (3:1 sulfuric acid: hydrogen peroxide) for 1 hour followed by thorough rinsing with ultrapure water (MilliQ, 18.2 MΩ resistance) and subsequent plasma cleaning for 15 minutes (Argon, PDC-002, Harrick Plasma). Lipid vesicles were prepared by extrusion through a 50-nm membrane (Whatman, Maidstone, UK) using an Avanti Mini-Extruder (Avanti Polar Lipids, Alabaster, AL). The vesicles consisted of 1-palmitoyl-2-oleoyl-sn-glycero-3-phosphocholine (POPC) with either 0 or 5.0 wt% of 1,2-di-(9Z-octadecenoyl)-sn-glycero-3-[(N-(5-amino-1-carboxypentyl)imino-diaceticacid)succinyl] (nickel salt) (18:1 DGS-NTA(Ni), both Avanti Polar Lipids, Alabaster, USA). POPC vesicles containing 0.01 wt% Oregon Green 488 1,2-dihexadeca-noylsn-glycero-3-phosphoethanolamine (OG-DHPE) were also used for some of the experiments in this work. The buffer solution used in the experiments was HBS buffer (10mM 2-[4-(2-hydroxyethyl)piperazin-1-yl]ethanesulfonic acid (HEPES), 150mM NaCl), adjusted to a pH of 7.4, filtered through a 0.2 μm membrane (AnaChem, Luton, UK) before use. Press-to-seal silicone isolators (4.5mm diameter, Grace Bio-Labs, Bend, Oregon, USA) were pressed on the cleaned cover slips, following the manufacturer’s instructions, to form SLBs within the silicon wells of the isolator by adsorption and subsequent rupture of the lipid vesicles prepared by placing a 20 μl drop of the vesicle solution (1 mg/mL lipids) on the glass surface. After the formation of an SLB from the lipid vesicle suspension (~30min) the vesicle solution was first replaced with buffer solution and then with a solution containing 0.25μg/mL of rCD2-6His-spacer-6His. For some experiments, the rCD2 molecules were labeled with Alexa Fluor 488 or 647 (Molecular Probes, Invitrogen, UK) *via* surfaced exposed lysines. Ni_2_+-NTA lipids were incubated with rCD2 in the SLB for 60 minutes after which the binding of proteins had reached equilibrium.

#### Sample preparation for imaging (TIRFM experiments Fig.2 *B-D* and 4)

Before imaging, approximately 10^6^ cells were resuspended in PBS (phosphate buffered saline, pH 7.4) and incubated in a microcentrifuge tube with the desired antibody fragments (as indicated in the text) for 30 min at room temperature (22°C). After the incubation step, the cells were washed three times with PBS by centrifugation and resuspension of the pellet (600×g, 2 min) and transferred to an SLB for imaging. After the slides were transferred to the microscope stage, cells were added and imaging was carried out within the first minutes following cell attachment room temperature.

#### Sample preparation for imaging (DHPSF experiments, Fig. 2 *E*)

Jurkat T CD48^+^ cells were labelled with Alexa 647-CD45 antibody (Gap8.3, anti-CD45) as previously described before being fixed in 4% paraformaldehyde (Sigma-Aldrich) and 0.2% glutaraldehyde (Sigma-Aldrich) for 60 minutes at room temperature. The fixed cells were washed three times in PBS and suspended in GLOX STORM buffer (PBS supplemented with 50mg/ml glucose (Sigma-Aldrich), 0.02-0.05 mg/ml catalase (Sigma-Aldrich), 0.8 mg/ml glucose oxidase (Sigma-Aldrich) and 7mg/ml MEA (Sigma-Aldrich). Coverslips (24 × 50 mm borosilicate, thickness No. 1, Brand) were coated with poly-L-lysine (Sigma-Aldrich, Mol wt 150-300 kDa) for 10 minutes, before a 1:100 dilution of 100 nm gold nanoparticles (Sigma-Aldrich) was added for 2 minutes. Coverslips were washed three times with PBS and 50 μL of the fixed cells were placed onto the coated coverslips. Fixed T cells were imaged for 200,000 frames with continuous 640 nm and 405 nm HiLo laser excitation and a 30 ms exposure (Tokunaga, et al., 2008). After reconstruction, a rolling-mean of the fiducial marker’s position over 50 frames was used to correct for drift in x, y and z.

#### Image analysis (TIRFM experiments)

Image analysis was performed using a combination of manual analysis (ImageJ, U. S. National Institutes of Health, Bethesda, Maryland, USA, http://imagej.nih.gov/ij/)) and custom-written MATLAB code (MATLAB R 2014b, The MathWorks, Natick, US).

##### i. Image acquisition and analysis (single-molecule tracking experiments for close-contacts on SLBs, Fig. 2 B-D)

Videos were obtained at a frames rate of 28.6 frames per second (exposure time: 33ms) simultaneously visualizing the distribution of wild-type CD45 (labeled with high FAB concentrations as described above) in one channel and trajectories of either wtCD45, HA-CD45-Halo, (mm)Lck-Halo or TCR-β-Halo in the second channel. Videos were analyzed using custom-written software (MATLAB). An interactive user interface allowed the user to select boundaries for each cell based on bright field images acquired during the data collection phase. Binary masks of the CD45 distribution for the selected cells were obtained by an average Z-projection over 200 frames in the CD45 channel, and applying an intensity threshold to the image (pixels were assigned a value of 1 if their intensity values I fulfilled the condition I > (mean(I_region_) −2 * std(I_region_)). The videos for the second channel (recording the trajectories of single molecule in the close-contacts) were analysed using the custom-written software (MATLAB) described in Weimann et al. The positions of the trajectories obtained were then mapped onto binary representations of the contacts created before using a custom-written MATLAB routine. Using these binary images, the routine also sorted the trajectories into two categories (“confined within” and “outside” the contact). The diffusion coefficients for those two categories were trajectories were extracted as described elsewhere (Weimann et al. 2013). The number of classified trajectories for a given cell was normalized by the corresponding mask area, *i.e.* the area “inside” and “outside” the contact, respectively.

##### ii. Image acquisition and analysis (simultaneous imaging of CD45 distribution in the close-contacts on SLBs and Ca^2+^ signaling / Fluo-4 fluorescence increase with TIRFM, Fig. 3 D; 4 D-F)

Data (videos) were acquired on the TIRF microscope described in section Total internal reflection microscopy at 37°C, using the 488 and 633 lasers for excitation and the corresponding dicroics and emission filters. Data were acquired using alternating excitation and time-lapse acquisition, with an exposure time of 100 ms, and a time between frames of 2 s. The videos were analyzed using custom-written MATLAB code. Briefly, cell positions were manually traced in the last frame of the Fluo-4 channel, and for each cell its mean intensity trace across the entire image sequence was calculated. The time of landing (t_land_) and Ca^2+^ release (t_Ca_) were determined in a semiautomated fashion from these intensity traces; tland was determined manually from the bright field image and tCa automatically by finding peaks in the derivative of the intensity trace. The time taken to trigger defined as t_Ca_ – t_land_. For the interval t_land_ to t_Ca_, the area of cell-surface contact was calculated from the CD45 channel for each cell by manually tracing the area inside the close contact outlined by CD45 fluorescence in a program-integrated GUI. From this analysis, we obtained the change of cell-surface contact area for individual cells until the time point of Ca^2+^ release t_Ca_. All further statistical analysis and fits of the traces were performed with Origin (OriginPro 9.1 G, OriginLab Corporation, Northampton, USA).

##### iii. Super-resolution microscopy of CD45 distribution in Jurkat T cells (Fig. 2 F,G)

CD45 was a labeled with Gap8.3 conjugated to the fluorescent dye Atto655 and made to emit intermittently by the addition of ascorbic acid (100μM; for details see Vogelsang et al., 2009). Movies of isolated fluorescent puncta could thus be obtained whose center positions were extracted using the software PeakFit (www.sussex.ac.uk/gdsc/intranet/microscopy/imagej/smlm_plugins), an ImageJ plug-in for superresolution analysis. Briefly, local maxima in each frame were fitted with a 2D-Gaussian described by seven parameters (position on two axes, standard deviation on two perpendicular axes and angle to the horizontal axis, amplitude, and offset). Finally, each single-molecule position was replotted using a custom macro written in ImageJ (http://rsb.info.nih.gov/ij/) as a 2D Gaussian profile defined by the measured integrated intensity and a width given by the average statistical error in localization of the centre (95% confidence interval, averaged over all single-molecule localizations; for further details see (Ptacin et al., 2010).

##### ib. DHPSF experiments (Fig. 2 E)

All DHPSF data was analysed by easy-DHPSF software (Lew et al., 2015) and was rendered using ViSP (Beheiry et al., 2013).

### Quantitative Treatment of Signaling

This model seeks to mathematically describe TCR dwell time inside close contacts. Close contacts are defined as any cell area from which CD45 is excluded; corresponding to close contacts described in Chang *et al.* and this study). Our experiments find that these contacts are initially small and grow over time (see Movie S3; Chang et al., 2016, Razvag et al., 2018). Cell-cell contacts appear to be highly dynamic in the absence of signaling. Since we are interested in signal initiation, we assume that close contacts form for finite periods (referred to as the contact duration).

In contrast to other models, the kinetic segregation (KS) model proposes that receptor triggering requires only that the TCR stays accessible to kinases within close contacts, protected from phosphatases that would otherwise terminate signaling, and for TCR phosphorylation to be sufficiently long-lived for downstream effects to be initiated. pMHC ligands, *via* trapping effects, serve only to increase the residence time of the TCR inside the close contact (Davis and van der Merwe, 1998; Davis et al. 2006). We therefore assume for our modeling (1) that when a close contact is formed, TCRs can diffuse in and out of the contact, and (2) that if the TCR is bound to a ligand inside the close contact, it is unable to leave. Any TCR that remains in the close contact for longer than 2 seconds, irrespective of its binding status, is assumed to be triggered. We chose t_min_ = 2 seconds, given that k_cat_(Lck) = 3.41 s^−1^ and that at a CD45/Lck ratio of 2:1, ITAM phosphorylation appears to be reduced by around 50%(18). We note that regulatory phosphorylation of Lck, that CD45 also counteracts, has very little effect on kcat and is therefore unlikely to affect this parameter, unphosphorylated and doubly phosphorylated Lck have similar k^−^ _cat_ (3.41 v 3.29). If we therefore assume that that effective ITAM phosphorylation is k_cateff_ «2, this still means that in 2 seconds 4 pTyr events would occur, which correspond to two pTyr ITAM signaling domains and which we assume is sufficient to initiate downstream signaling. Furthermore, there is experimental evidence that TCR triggering occurs within this time-frame upon pMHC binding(10, 22–25). In essence, by introducing t_min_ we consider a distribution of phosphorylation times with a sharp lower boundary created by the maximal turnover rate of Lck in case of immediate substrate binding (*i.e.* k_cat_). If a TCR leaves but re-enters shortly thereafter, then that TCRs sojourn inside the close contact is considered separately with a restarted clock, assuming that triggering occurs after a TCR has occupied the close contact continuously for 2 seconds and not cumulatively over multiple excursions. The model calculates TCR density across the close contact based on the rate of TCR entry into this area (for a given contact radius, initial TCR density and TCR diffusion coefficient). Therefore, it can also account for changes in TCR entrance rate and in the dwell-time of TCRs already present inside the close contact as it grows. To achieve this higher level of accuracy, this model requires moving-boundary coupled partial differential equations that are computationally expensive. While the problem is solved numerically on a disc, the solution is fully two dimensional and not radially symmetric (although the domain is). Our simulation method is based on a finite element discretization implemented in MATLAB.

#### Modelling of receptor triggering using moving-boundary coupled partial differential equations to account for close contact growth

See Figs 1–5, Supplementary table 2 and Movie S1

##### a. Model formulation

Since close contact zones (CCZs) grow on time-scales similar to the diffusion of the TCR, changes in TCR density in CCZs need to be described by a coupled system of moving-boundary partial differential equations (PDEs),

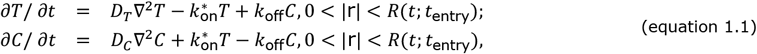

where *T*(**r**,*t;t*_entry_) and *C*(**r**,*t;t*_entry_) represent free and ligand-complexed TCRs diffusing with coefficient D_T_ and D_C_, respectively, and with the receptors undergoing reversible binding with first-order rates 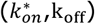. Note that 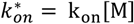 where k_on_ is the bimolecular on-rate (in units of *μm*^2^/*s*) and [M] is the ligand concentration (in units of *μm*^−2^). We have assumed that TCRs within a CCZ do not compete for pMHC and this is reflected in using first-order kinetics for binding. This approximation is reasonable when the number of pMHC is larger than the number of bound TCRs at all times within the CCZ. The boundary conditions for the disc domain of radius *R* are adsorbing for *T* and no flux for *C*,

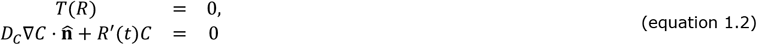

Importantly, the domain area grows linearly in time and therefore,

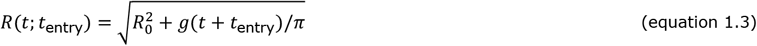

where *g* is the growth rate (in units of *μm*^2^/*s*) and *t* is time. The initial conditions at *t* = *t_entry_* are as follows,

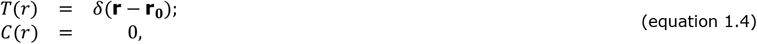

where r_0_ = (R_0_ -ε,θ). The additional term *R′(t)C*, which reflects the rate of growth in the region, is a necessary addition to the usual Neumann condition in order to prevent mass of *C* leaving the domain. To see this, consider the change in total mass 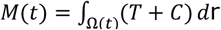,

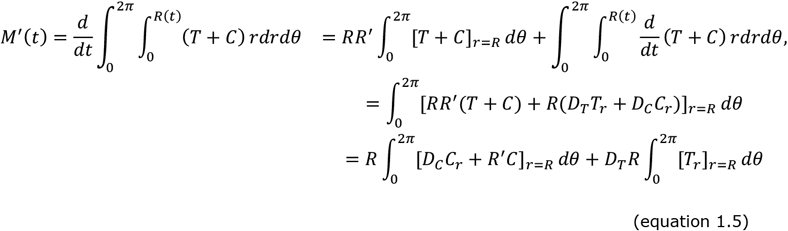

The flux of T-cell receptors in complex (*C*) through the boundary should be zero which gives rise to the boundary condition. In the case of modeling CCZs of fixed size (Fig. 3B-E), *g* = 0 in equation 1.3. The parameter *ϵ* is the distance from the boundary that the TCR is initialized in its exploration of the CCZ. For technical reasons, the initial location cannot be exactly on the boundary as one would ideally like since the mathematical formulation of the dwell time would not be well posed. This is because a Brownian walker initially on the boundary will interact with the boundary an infinite number of times. Therefore, we must start the particle just inside the domain and we have used a value of *ϵ* = 0.09 for the numerical simulations.

##### b. Model output

The output of the model is the probability (*P_s_*) that a single receptor has remained within the CCZ for more than 2 seconds, for contact duration (*t_f_*),

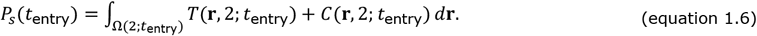

The time-dependent rate of TCR entry into the domain (*k_t_*(*t*)) is expected to be proportional to the size of the domain, which increases over time. Using previously derived results (see Equations 11 in Weaver, 1983), we find,

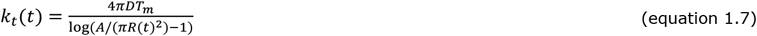

where *A* = 415 *μm*^2^ is the cell surface area, *T_m_* = 100 *μm*^−2^ (varied over the simulations; see also Table S3) is the TCR density far away from the CCZ, and *D_τ_* = 0.05*μm^2^/s* (varied over the simulations, see also Table S3) is the TCR diffusion coefficient. With these numbers, we find that the rate of TCR entry into the domain (*k_t_*) increases from ≈ 4 *s*^−1^ to ≈ 18 *s*^−1^ as the domain radius increases from 0.01 μm to 2 μm. Note that since we numerically solve the problem by a finite element method, the coordinate singularity at r = 0 does not need to be resolved in this setting.

Given that multiple receptors can enter the CCZ during the contact duration (*t_f_*), we need to calculate the probability that at least one TCR has remained within the domain for more than 2 s (*P_m_*, referred to as “triggering probability”). The number of TCRs that have entered the domain in time interval [*t_i_,t_i_* + Δ*t*] can be estimated as *k_t_*(*t_i_*)Δ*t* so that *P_m_* is estimated as follows,

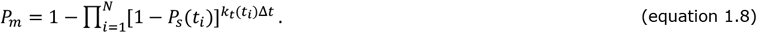

In the case where *P_s_* and *k_t_* are constants:

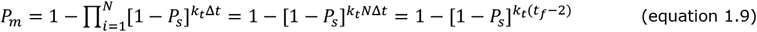

When considering the limiting case of kt(t) = 0 for the growing CCZ model, we have assumed a small initial radius (0.01 μm) and therefore assumed that the CCZ is empty of TCRs. In cases where we do not have a growing CCZ (*g* = 0, Fig. 3C), a term is included in the expectation representing the initial number of TCRs in the CCZ at *t* = 0,

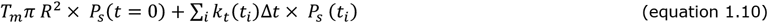

All plots for the theoretical modelling of TCR triggering were generated in MATLAB (MATLAB R 2014b, The MathWorks, Natick, US) using the equations derived in this section.

#### Quantification and Statistical analysis

Data were analysed by Graphpad Prism and Origin Lab built-in T-test (unpaired, two tailed), and results were considered significant when p < 0.05. Other statistical parameters including the number of replicates, fold-changes, percentages, SEM, SD, number of cells, number of tracks and statistical significance are reported in the Figures, Figure legends and supplementary tables.

#### Data and Code Availability Statement

The data sets generated during and analyzed during the current study as well as all custom-written software are available from the corresponding author on reasonable request.

## II. Supplementary Tables

**Supplementary Table 1.** Characterization of molecular behavior in close contacts using single-molecule fluorescence microscopy. Values from the data presented in Fig. 2. The automated procedure used to dissect the images and define the cell-SLB contacts are described in the Online Methods.

| | # Cells/<br>contacts | Average<br>contact<br>area<br>[$\mu\text{m}^2$] | Average<br>contact<br>radius<br>[$\mu\text{m}$] | Confined*/<br>total<br>molecules | Confined<br>molecule<br>density /<br>total<br>density† | Diffusion coefficient<br>(mean±s.d. of<br>distribution measured;<br>$\mu\text{m}^2/\text{s}$) | |
| --- | --- | --- | --- | --- | --- | --- | --- |
|  |  |  |  |  |  | Inside<br>of contact | Outside<br>of contact |
| **CD45** | 39 | 25±20 | 2.8±2.6 | 65/969 | 0.13±0.03 | | 0.113<br>±0.179 |
| **Lck** | 16 | 16±21 | 2.2±2.6 | 234/1527 | 0.56±0.07 | 0.151<br>±0.294 | 0.104<br>±0.185 |
| **TCR** | 16 | 13±12 | 2.1±2.0 | 48/375 | 0.40±0.06 | 0.054<br>±0.107 | 0.050<br>±0.074 |

\* Trajectories were classified as “confined” if they could be mapped exclusively onto the inner area of a contact (denoted by “1” in the binary representation of the contacts, cf. Fig. **1 b-d**, *bottom panels*).
† Quoted is the fraction of a molecule’s density “in” the contact over its total density in the visible area of the cell.

**Supplementary Table 2.** Parameters used in the calculations shown in Figs 3, 4, 5 and 6.

|  | Fig. 3A | Fig. 3B | Fig. 3C | Fig. 4A | Fig. 4B | Fig. 4C | Fig. 5A | Fig. 5B | Fig. 5C | Fig. 5D | Fig. 6A | Fig. S6B |
| --- | --- | --- | --- | --- | --- | --- | --- | --- | --- | --- | --- | --- |
| Cell surface area<br>[$\mu\text{m}^2$] | 415<sup>3</sup> | | | | | | | | | | | |
| Number of close<br>contacts | 1 |  |  |  |  |  |  |  |  |  |  |  |
| Close contact radius<br>[$\mu\text{m}$] | *variable*<sup>1</sup> | | | | | | 0.22<sup>3</sup>/<br>0.44<sup>3</sup> | 0.22<sup>3</sup> | | *variable* | 0.22<sup>3</sup> | |
| # TCR copies/cell | 41,500<sup>2</sup> |  |  |  |  |  |  |  |  |  |  |  |
| # CD45 copies/cell | 200,000<sup>2</sup> |  |  |  |  |  |  |  |  |  |  |  |
| # Lck copies/cell | 40,000<sup>2</sup> |  |  |  |  |  |  |  |  |  |  |  |
| Fraction of TCR<br>segregation | 0.63<sup>2</sup> |  |  |  |  |  |  |  |  |  |  |  |
| CD45:Lck ratio | 2.5:1 |  |  |  |  |  |  |  |  |  |  |  |
| TCR diffusion [$\mu\text{m}^2/\text{s}$] | 0.05<sup>2</sup> | | | | | | | | | | | |
| $k_{\text{on}}$ (pMHC) | NA | | | | | | | | 0.1 | 0.1 | 0.1 | *variable* |
| Agonist pMHC<br>[molecules/$\mu\text{m}^2$] / $k_{\text{off}}$ | NA | | | | | | | | 30 / 1 | 30 / 1 | *variable* | |
| Non-agonist pMHC<br>[molecules/$\mu\text{m}^2$] / $k_{\text{off}}$ | NA | | | | | | | | 300 / 50 | 300 / 50 | *variable* | NA |
| $t_{\text{min}}$ [s] | NA | 2<sup>3</sup> | | *variable* | 2<sup>3</sup> | | 2<sup>3</sup> | *variable* | 2<sup>3</sup> | | | |
| Triggering<br>Probability: residence<br>time > $t_{\text{min}}$ | NA | *result* | NA | *result* | | | | | | | | |
| T cell-APC contact<br>duration | 120 s |  |  |  |  |  | *variable* |  |  | 120 s |  |  |

<sup>1</sup> Growth rate, $g$ [ $\mu\text{m}/\text{s}$ ];
<sup>2</sup> Experimentally determined for CD4 T cells isolated from PBMCs;
<sup>3</sup> Referenced from literature, *c.f.* main text for details;

**Supplementary Table 3.** Detailed pMHC/TCR density and kinetic parameters used for the calculations shown in Fig. 6 B.

| For CD8+ T cells |  |  |  |  |  |  |  |  |
| --- | --- | --- | --- | --- | --- | --- | --- | --- |
| http://www.nature.com/nature/journal/v464/n7290/full/nature08944.html |  |  |  |  |  |  |  |  |
| Peptide | Potency | $A_c * k_{on}$ [$\mu\text{m}^4/\text{s}$] | $k_{on}^1$ [$\mu\text{m}^2/\text{s}$] | $k_{off}$ [$\text{s}^{-1}$] | pMHC<br>[molecules<br>/$\mu\text{m}^2$] | EC<sub>50</sub> [$\mu\text{M}$] for peptide<br>(IL2 production) | P<sub>s</sub> (eq 1.6) | Triggering<br>probability P<sub>m</sub> (eq<br>1.9) |
| **OVA** | Agonist | 0.012 | 1.2 | 10.8 | 30 | 3.00E+10 | 0.002523396 | 9.33E-01 |
| **A2** | Agonist | 0.0035 | 0.35 | 4.7 | 30 | 1.60E+09 | 0.004031146 | 9.86E-01 |
| **V-OVA** | Non-agonist | 0.000004 | 0.0004 | 1.4 | 30 | 1.10E+05 | 0.000468371 | 4.94E-02 |
| **R4** | non-agonist | 0.0000044 | 0.00044 | 2.6 | 30 | 2.50E+05 | 7.98536E-05 | 9.29E-03 |
| **G4** | weak agonist | 0.000027 | 0.0027 | 1.3 | 30 | 1.95E+06 | 0.00037106 | 3.28E-01 |
| **E1** | weak agonist | 0.000013 | 0.0013 | 2.4 | 30 | 2.20E+05 | 3.15887E-05 | 3.33E-02 |

| For CD4+ T cells |  |  |  |  |  |  |  |  |
| --- | --- | --- | --- | --- | --- | --- | --- | --- |
| http://www.jimmunol.org/content/195/8/3557.long |  |  |  |  |  |  |  |  |
| Peptide | Potency | $A_c * k_{on}$ [$\mu\text{m}^4/\text{s}$] | $k_{on}^1$ [$\mu\text{m}^2/\text{s}$] | $k_{off}$ [$\text{s}^{-1}$] | pMHC<br>[molecules<br>/$\mu\text{m}^2$] | EC<sub>40</sub> [$\mu\text{M}$] for peptide<br>(IL2 production) | P<sub>s</sub> (eq 1.6) | Triggering<br>probability P<sub>m</sub> (eq<br>1.9) |
| **hb** | agonist | 0.000233 | 0.0233 | 1.22 | 30 | 0.0004 | 0.00380000000 | 9.83E-01 |
| **t72** | weak agonist | 0.000116 | 0.0116 | 1.53 | 30 | 0.023 | 0.00110000000 | 6.93E-01 |
| **i72** | non-agonist | 0.000145 | 0.0145 | 2.99 | 30 | 7 | 0.00016872000 | 1.66E-01 |
| **a72** | weak antagonist | 0.000129 | 0.0129 | 3.79 | 30 | 60 | 0.00005261600 | 5.49E-02 |

<sup>1</sup>The parameter $A_c$ stems from method used to measure the rates (*i.e.* monitoring abrupt decrease/resumption in thermal fluctuations of a biomembrane force probe) which do not measure $k_{on}$ directly. We assumed $A_c = 0.01$ (*i.e.* 1% of 1 $\mu\text{m}^2$ ) for calculating $k_{on}$ based on the author's estimate in Huang *et al.*, Nature, 2010 (11).

## III. Supplementary Figures

**Supplementary Fig. 1.**
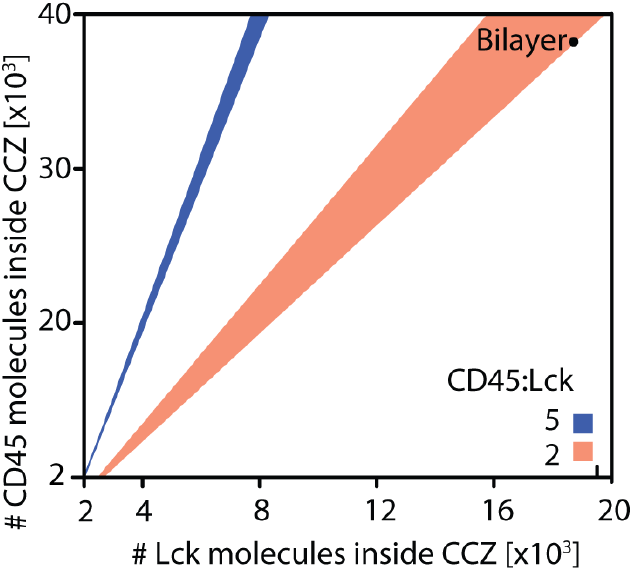
Quantitative analysis of CD45 and Lck density. Graphical depiction of the CD45 to Lck ratio of 5 (blue; corresponding to the average number of CD45 and Lck found at the cell surface) or 2 (orange) for varying numbers of total CD45 and Lck molecules in the T cell membrane (±0.25). Considering the single-molecule quantification of CD45 and Lck found inside the close contacts (see Fig. 1) the ratio of CD45 to Lck in the bilayer is close to 2.2.

**Supplementary Fig. 2.**
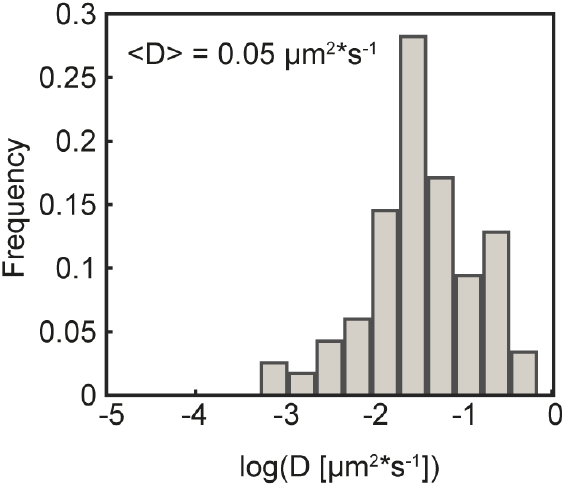
TCR diffusion at close-contacts between T cells and support lipid bilayers by single-particle tracking. Histogram of diffusion coefficients D obtained from linear fits (5 points, interval 35ms) to mean-square displacement (MSD) of single TCRs (data presented in **Fig. 2d**). Trajectories for CD45 and Lck were analyzed analogously to calculate their respective diffusion coefficients; the diffusion coefficient quoted in the **Supplementary table 1** is the mean (<D>) and the error the s.d. of the distribution of D.

**Supplementary Fig. 3.**
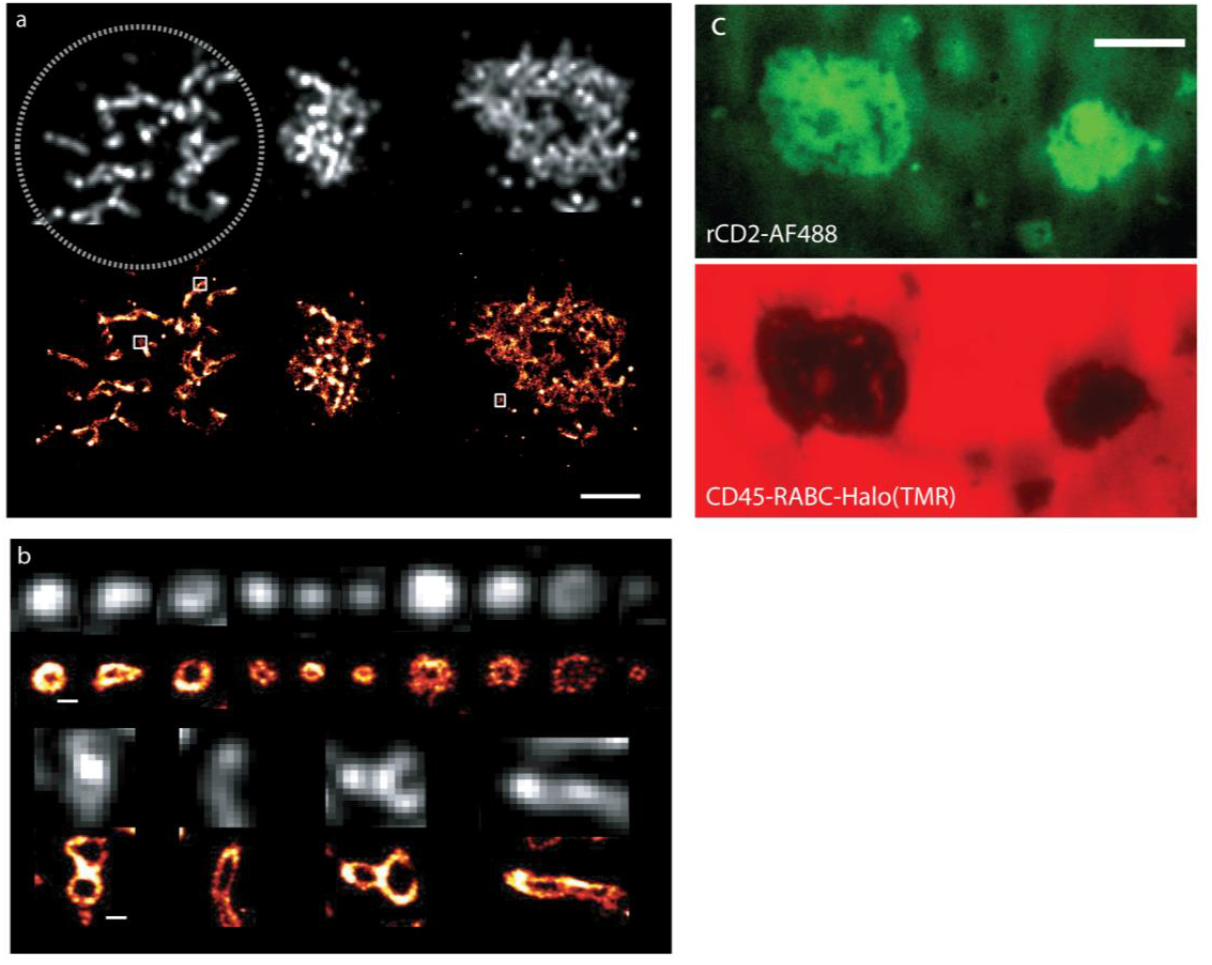
CD45 is equally distributed on resting T cell surfaces and segregates from close-contacts between T cells and glass surfaces at sub-μm length scales. **(a,b)** 2D super-resolution *d*STORM image of CD45 in early T cell contacts formed with IgG coated glass surface; average (*top*) and reconstructed image (*bottom*, scale bar 3 μm). CD45 is excluded from contacts as small as 78 nm in diameter. Representative field of view (**a**) and (**b**) galley of 20 individual close-contacts sampled across > 5 cells (scale bar 100nm). Note that the CD45 distribution appears to be homogeneous across these contacts if the resolution is limited by diffraction. (**c**) The spatial organization of rCD2 (Alexa Fluor 488- tagged, green, 1000 molecules/μm^2^, top) and CD45-RABC-Halo (TMR labeled, red, bottom) in contacts of CD48+ Jurkat T-cells and SLBs containing rCD2 and CD45-RABC-Halo (4:1) 8 minutes after cells landing on the SLB.

**Supplementary Fig. 4.**
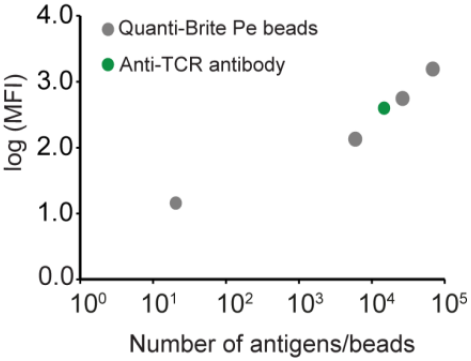
Determination of total contact area at time of Ca^2+^ release for close-contacts between T cells and rCD2-SLBs and measurement of total TCR numbers in the cell line used for experiments. Quantification of the number of TCR per cell measuring mean fluorescence intensity of anti-TCR PE labeled antibody by FACS and QuantiBrite PE beads.

**Supplementary Fig. 5.**
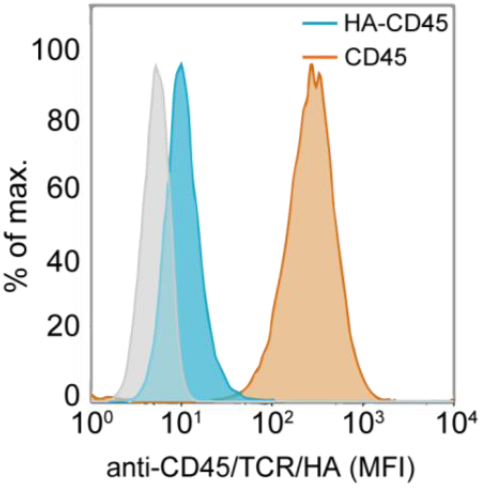
Measurement of HA-CD45 expression levels. Mean fluorescence intensity of anti-HA (clone HA-7, Sigma, HA-7) or anti-CD45 (Gap 8.3) antibodies on Jurkat T cells transduced with HA-CD45. In grey, anti-HA staining of untransduced Jurkat T cells.

**Supplementary Fig. 6.**
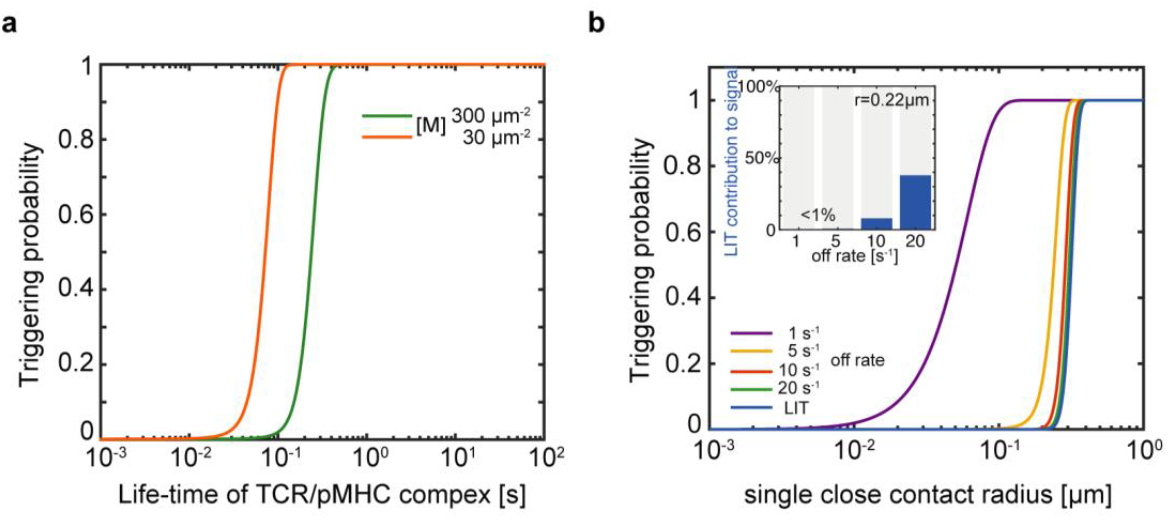
The effect of TCR/pMHC complex life time on triggering probabilities and the contribution of LIT to the overall signalling probability at varying close-contact radii and for ligands with different k_off_ rates. (**a**) Triggering probability as a function of pMHC-TCR life-time (1/k_off_) for different pMHC densities [M]. (**b**) Dependence of the triggering probability for TCRs on contact radius in the presence of agonist pMHC with varying k_off_ (30 pMHC molecules/μm^2^) and in the absence of pMHCs (LIT; blue line). The insert shows the extent to which triggering results from TCRs which are triggered without binding to pMHC (‘LIT’).

## Supplementary Movies (descriptions)

**Supplementary Movie 1**

**Animation of the changes in the TCR probability density across a growing close contact over time corresponding to the modelling shown in Fig. 1c**. Probability of occupation is plotted on the z-axis, the ‘time’ given in the title is the time post initial contact and the ‘remaining mass’ refers to the probability that the TCR is still found within the close contact. Close contact growth rate is set to g = 0.1 μm^2^/s.

**Supplementary Movie 2**

**CD45-RABC spontaneously segregates from CD2-mediated close-contacts formed by Jurkat T-cells (CD48+) with CD2- and CD45RABC-containing SLBs.**

Representative movie showing simultaneous rCD2 accumulation and segregation of CD45RABC-Halo from stable cell-bilayer contacts of CD48+ Jurkat T-cells interacting with a rCD2- and CD45RABC-Halo containing SLB (rCD2:CD45RABC-Halo 4:1). The movie combines raw data for the CD2 channel (*i.e.* SLB contains Alexa Fluor 488-tagged CD2 (green, left) with a simultaneously-acquired movie of the CD45RABC-Halo channel (TMR, red, right). The movie is played back 10-fold faster than real-time (5 frames per second). Scale bar, 5 μm.

**Supplementary Movie 3**

**Ca^2+^ release as measured by change in Fluo-4 fluorescence in Jurkat T-cells forming contacts depleted of CD45 (labeled) with CD2-containing SLBs**. Movie collage for a CD48+ Jurkat T-cell; the movie combines raw data for the CD45 channel *(i.e.* CD45 labeled with Alexa Fluor 568-tagged Gap 8.3 Fab *(red, left)* with a movie of the Fluo-4 channel *(green, right).* The movie is played back 10-fold faster than real-time (5 frames per second). Scale bar, 5 μm.

